# Refinement of Draft Genome Assemblies of Pigeonpea (*Cajanus cajan*)

**DOI:** 10.1101/2020.08.10.243949

**Authors:** Soma Marla, Pallavi Mishra, Ranjeet Maurya, Mohar Singh, D. P. Wankhede, Anil. K. Gupta, N. S. Rao, S. K. Singh, Rajesh Kumar

**Author notes:** Soma Marla, Pallavi Mishra, Ranjeet Maurya contributed equally.

## Abstract

Genome assembly of short reads from large plant genomes remains a challenge in computational biology despite major developments in Next Generation sequencing. Of late multiple draft assemblies of plant genomes are reported in many organisms. The draft assemblies of *Cajanus cajan* are with different levels of genome completeness; contain large number of repeats, gaps and segmental duplications. Draft assemblies with portions of genome missing, are shorter than the referenced original genome. These assemblies come with low map accuracy affecting further functional annotation and prediction of gene component as desired by crop researchers. Genome coverage *i.e.* number of sequenced raw reads mapped on to certain locations of the genome is an important quality indicator of completeness and assembly quality in draft assemblies. Present work was aimed at improvement of coverage in reported *de novo* sequenced draft genomes (GCA_000340665.1 and GCA_000230855.2) of Pigeonpea, a legume widely cultivated in India. The two assemblies comprised 72% and 75% of estimated coverage of genome respectively. We employed assembly reconciliation approach to compare draft assemblies and merged them to generate a high quality near complete assembly with enhanced contiguity. Finished assembly has reduced number of gaps than reported in draft assemblies and improved genome coverage of 82.4%. Quality of the finished assembly was evaluated using various quality metrics and for presence of specific trait related functional genes. Employed pair-end and mate-pair local library data sets enabled to resolve gaps, repeats and other sequence errors yielding lengthier scaffolds compared to two draft assemblies. We report prediction of putative host resistance genes from improved sequence against *Fusarium* wilt disease and evaluated them in both wet laboratory and field phenotypic conditions.

## Introduction

Rapid developments in sequencing technologies facilitated generation of several draft assemblies in plants. These are valuable resources for elucidating genetic information and understanding biology of the crop. However, each of these draft assemblies have strengths and weaknesses as were sequenced and assembled based on different algorithms [1,2]. Draft assemblies differ on the sequencing technology and also the assembly software employed. One assembly may be conservative in selection of reads resulting in low genome coverage but with many gaps. Another assembler is vigorous, yielding more contigs but with many errors. Draft genomes are typically sets of large contingent of assembled contigs and scaffolds that are often fragmented due to presence several gaps interlaced by repetitive regions. In a misassembly different contigs are improperly joined. Mis-joins problem arises due to inversions, relocation or a translocation. Gaps arise also due to incorrect insertion or deletion of a sequenced read in a misassembly. An inversion or a translocation alters placement of a contig on to scaffold belonging to a different chromosome. Hence, annotation of unfinished and partially assembled genomes creates ambiguities while accessing complete genetic information as desired by biologists.

In misassemblies some of the reasons for incompleteness include 1.Gaps appearing due to polymorphisms in complex genomes where reads on either side of a gap representing two haplotypes belonging to two separate chromosomes, 2. Abundance of repeat elements, multiple ways to fill the gaps and confusing the assembler thus leaving a gap unfiled, 3. Lack of more reads to cover that part of the genome, requiring additional library of reads to fill the gaps. Besides, in draft genome assembly base call errors, variations in read depth coverage also cause gaps and pose serious computational challenges while connecting nodes in a *De Bruijn* graph [3].

Complex eukaryotic genomes are known to contain large volume of near identical copies of DNA repeats and fragments. Various types of repeats present in genomes of wheat, pigeonpea, maize or potato include transposable elements, highly conserved gene clusters and segmental duplications. Presence of identical (or near identical) DNA fragments further complicate computational assembly. During pre-assembly, short reads of equal size tend to be masked together and complicate construction of *De Bruijn* graphs [4]. Recently introduced third generation single molecule real time technologies [5] and Oxford Nano pore technologies [6] generate large sized reads which can readily be inserted for filling gaps caused by repetitive elements. However, due to low level of sensitivity, high sequencing error rates and expensive technologies many plant researchers are opting to short read sequencing technologies. Two draft *de novo* genomes compared in the present study are short read assemblies generated from second generation sequencing technologies. Abundance of repeats obviates gap closing and responsible for low levels of genome coverage reported in draft assemblies. Along with reads, modern sequencing platforms generate paired end reads or mate- pairs. The mate pair libraries are generated in different sizes (ranging from 3bp to 5bp) and orientations. Hence they could serve as potential inserts while filling gaps. Mate pair libraries are recommended as a potential approach to mitigate repeats in computational assembly. In the present work we demonstrated incorporation of suitable mate pairs to metassembly for gap closing, which in turn yielded significant improvement of both genome coverage and quality of the finished Pigeonpea assembly.

Major techniques suggested for gaining contiguity and higher coverage in draft genomes broadly include, use of long inserts for gap filling [7] assembly reconciliation, hybrid assembly [8], filtering repeats [9] and iterative mapping using short reads to close the remaining gaps [10]. Use of paired end or mate pairs for filling the gaps is a robust computational approach [11]. Reconciliation approach for closing gaps and correcting misassemblies involves comparing available data sets from different draft genomes of same or related species, mapping their reads and finally merge them together to gain improved scaffold lengths with higher contiguity [12].

Pigeonpea (*Cajanus cajan (L)* Millsp. *cv*. Asha) is a major food legume grown in India is diploid (2n = 22) with genome size of 833.07 Mbp [2]. Widely cultivated and is a major source of dietary proteins in India with annual production of 2.31mt and productivity of 678 kg/ha [13]. Prevailing low crop productivity may be attributed to absence of high yielding cultivated varieties possessing resistance to various pests and diseases. In plants, resistance genes (R genes) play important roles in recognition and protection from invading pests and pathogens. A few sources of resistance to biotic stresses can be found in available germplasm collections. Resistance genes are identified and found primarily organized in individual clusters that are strictly linked across the genome [14]. Modern plant breeding techniques such as Marker assisted and Genomic selections develop superior crop varieties making use of genomic resources and genetic information emitting from sequenced genome projects. Pigeonpea genome was *de novo* sequenced independently by [1,2]. These draft assemblies, available in public domain (GCA_000340665.1 and GCA_000230855.2) are valuable resources for breeders. However, both the assemblies are incomplete with sizable number of fragmented contigs and gaps. Lack of accurate genetic information is a major limitation towards prediction of gene compliment associated with desirable traits. Hence our primary objective in the present work is to generate a more contagious finished assembly with improved genome coverage. We report a finished assembly based on genome reconciliation approach that first compares the two available draft assemblies, scoring matching blocks at each location followed by their merger. Metassembler tool employed in the present study detected gaps and filled them iteratively using right sized inserts from local pair-end and mate-pair libraries. Completeness and map accuracy of the reconstructed assembly was verified for the presence of conserved plant resistance genes (R genes). Here we report prediction of putative R genes, their isolation and PCR screening of a set of known cultivars against *Fusarium* wilt disease in both laboratory and field conditions.

## Results

### Improvement of the draft genome assemblies employing reconciliation algorithm

Reconciliation assembly approach was employed in the present work to refine the fragmented draft genome assemblies A1 and A2. For selection of optimum K-mers, hybridSPAdes [15], was employed and combinations ranging from 21 to 55 were evaluated. We observed with k-mer sizes 21, 33, 55 and 77 yielded few fragmented sequences, less number contigs with high N50, mean and median scaffold lengths in superior assemblies. The Illumina HiSeq sequence reads resulted in 46,979 reads with the N50 length of 24,087. Metassembler was employed for merging of two assemblies. Metassembler implements reconciliation algorithm to refine and obtain reconstructed genome. In order to capture the suitable reference assembly set for alignment during merger process we examined the required order in which assemblies A1 and A2 are to be chosen as master set (GCA_000230855.2) and slave sets for aligning with the former, (GCA_000340665.1). We observed that choice of A1 as master set with and A2 as slave set resulted in a highly contiguous superior assembly. Superiority of resulting merged metassembly was systematically evaluated with the compression-expansion (CE) statistics. Gaps present in the scaffolds were closed using mate pairs. The remaining gaps were filled by searching unique contig end sequences against unused reads. We observed that repeat structure analysis and resulted significant reduction of gaps and contributed to prediction of specific genes. The improved assembly had 46,979 contigs with total size of 548.2 Mb and covers 82.4% of the genome with high contiguity (**Table 1**).

**Table 1:**
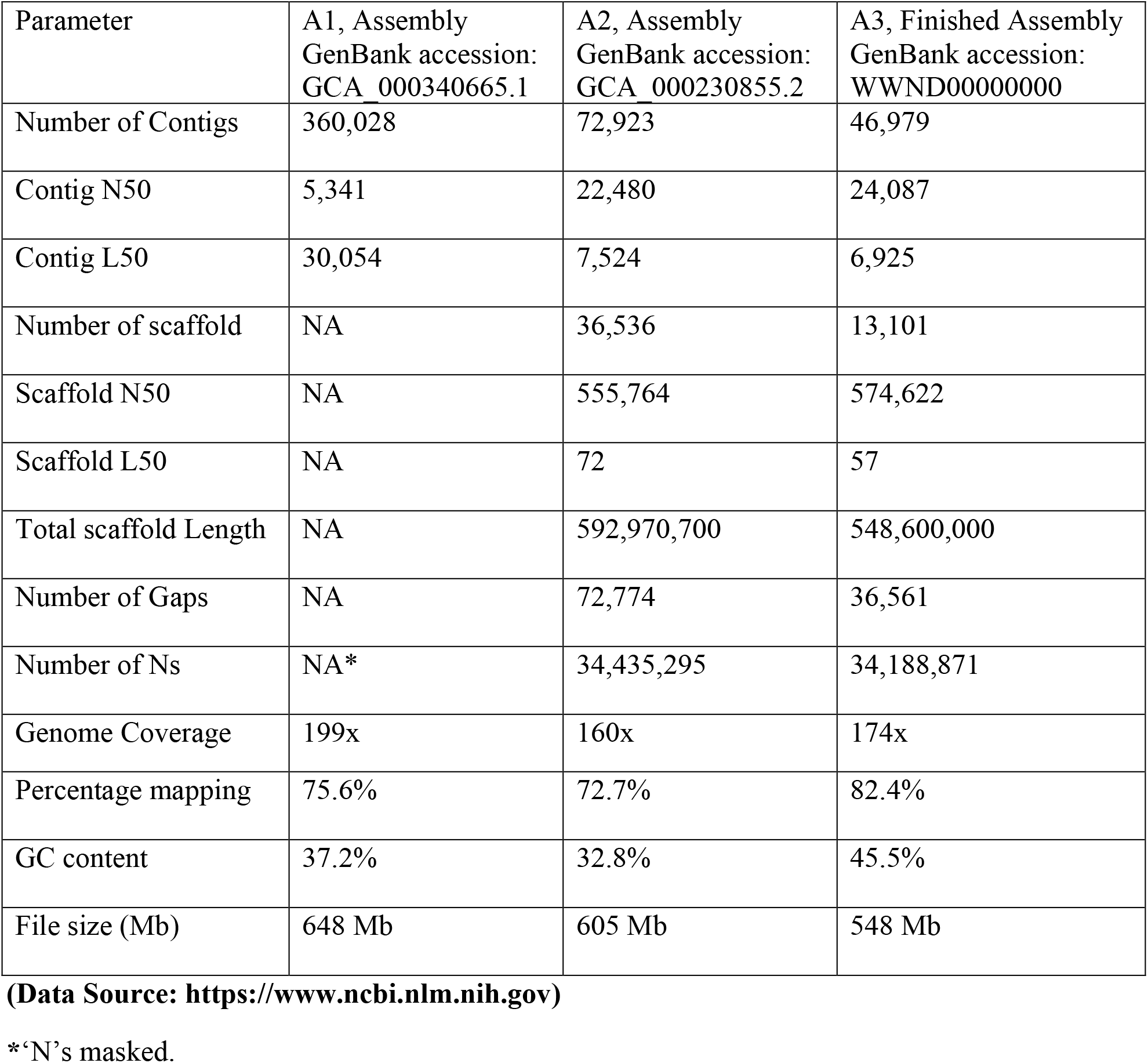
Genome assembly statistics of draft assemblies A1, A2 and finished A3 assembly.

### Read alignment/mapping of pigeonpea

Read mapping increased from 75.6% and 72.7% in two compared misassemblies to 82.4% in finished metassembly. More reads were found to be mapped to merged assembly than in A1 and A2 misassemblies. Mapping depth is a measure of number of reads used for aligning the finished genome. It also helps to estimate the extent of similarity between final finished assembly and the compared misassemblies. Among the two draft assemblies, A2 is superior to A1 in depth of read coverage. Relatively higher read depth in A2 misassembly can be attributed to the high-identity Illumina reads used both in initial assembly and later polishing steps. Our finished final assembly in terms of depth of coverage is superior to A2, with more gaps filled. In addition, refined assembly has more GC-rich regions (**Table 1)** and improved gene component predicted. The total GC content in A1 and A2 assemblies had GC content i.e. 37.2% and 32.8% respectively and enhanced to 45.5% in metassembly reported in the present work. The improvement in GC rich fraction and of N50 values in both contigs and scaffolds in the finished genome was achieved largely due to gap filling. High GC content is known to be associated with concentration of coded genes in certain regions of genome [16]. In the present study high GC content obtained in refined assembly A3 has contributed to increased number of predicted genes in the finished genome of pigeonpea.

### Metassembly, annotation and quality assessment

Two draft assemblies were merged and reassembled employing two approaches as described above. We wanted to ascertain which type of mate-pair libraries effectively resolve repeat problem. In assembly employing Meta assembler tool we used in one experiment only 648 Mb library and in the second 605 Mb and 548 Mb libraries taken together. Initially we used all the single paired read data sets available (minus two mate-pair data sets) of A1 along with all data sets from A2. In the second treatment included the two mate-pair data sets from A1 along with all full data available from A2. At the end of analysis, all the output values and statistical metrics were collected for comparative performance analysis. We observed that all the available Pigeonpea mate pair libraries taken together resulted improvement in genome coverage. It is presumed that incorporation of variable size mate pair inserts helped in gap closing during assembly.

In our final assembly the contig N50 is increased by 24,087, and scaffold N50 increased by 574,622. Total number of gaps decreased across the genome by 50.23%, comprising from A2 (**Table 1**). It is observed that the order in which the input draft assemblies are inputted to Metassembler drastically influences the alignment quality and the resulting read coverage. In primary assembly we treated Assembly A1 as master and aligned it with Assembly A2. In other variant we used Assembly A2 as master and aligned against its counterpart Assembly A1. Output of resulting primary assembly yielded us a scaffold length 548,600,000. We initially used unpaired reads for assembly adopting overlapping read approach. As no significant improvement was observed in both read mapping depth and eventual coverage we resorted to available mate paired libraries to close gaps. We used mate pairs during different alignment steps during metassembly and succeeded in resolving repeat problems.

### Closure of repeat-derived gaps

For each round of alignments undertaken between A1 and A2 misassemblies, metassembler builds a graph, with vertices being the above alignments and edges joining two alignments. If both have the same direction, they are readily rearranged in to a single block thus providing contiguity. In case, where the examined genomic segments from two misassemblies do not share same direction, indicates the existing distance from each other and need to fill the prevailing gaps. In such cases, variable sized local pair-end and mate-pair libraries could offer right inserts to fill these gaps. While building the graph, metassembler searches the mate-pair library for right sized inserts to complete the shortest path between any of these contigs, to fill a gap.

We evaluated the closure performances of the Gapcloser and Gapfiller tools on the repeat derived gaps using the raw mate pair reads. We first tested the performance of each tool using the raw mate pair reads. Both the above tools used first raw pair end and Mate pair libraries. We monitored the gap closure efficiency by evaluating number of gaps closed. In improved pigeonpea assembly, we estimated 37,145 repeat-derived gaps of which 584 gaps and 322,780 nucleotides out of total 34511651 were closed. The gap sizes ranged from 200 bp to 15,510 bp. Gap closer was more efficient by filling most of the gaps with 82.4% and with low error rates. We achieved improved contiguity by using long mate-pairs to fill gaps in assembly and there by achieving higher coverage in the finished assembly. Two-draft genome assemblies A1 (GCA_000340665.1) and A2 (GCA_000230855.2) are used in the present study to improve scaffold contiguity and achieve read coverage completeness of Pigeonpea genome. Draft assembly A1 had 360,028 contigs with N50 and L50 of 5,341 and 30,054 respectively. Reported genome coverage was 199x with a similarity of 75.6 %. Draft Assembly A2 had 72,923 contigs with N50 and L50 of 22,480 and 7,254 respectively. A2 had 592,970,700 scaffolds with reported genome coverage of 160x with a similarity of 72.7 %. We present an improved reference assembly of pigeonpea genome.

### Completeness of the merged assembly

The BUSCO [17] evaluation of completeness of the conserved proteins in all three assemblies of the pigeonpea genome sequence predicted that it was 94.02% complete in A3 assembly, where a proportion of total 1,440 BUSCO groups were searched, the genome assembly found to contain 1,321 complete single-copy (S) BUSCOs, 33 complete duplicated (D) BUSCOs, 57 fragmented (F) BUSCOs, and 29 missing (M) BUSCOs. Whereas comparatively in A1 and A2 assemblies it was found 85.27% (S:76.87%, D:8.40%, F:5.62%, M:9.09%) and 87.9% (S:80.9%, D:7%, F:5%, M:7.1%) complete respectively (**Supplementary Table 1**). The gene completeness as measured by BUSCO is increased in improved assembly, while the numbers of fragmented and missing BUSCO genes are reduced. This genome comparison can be used to help such draft assemblies towards becoming finished genome.

### Functional annotation of predicted gene content

FGENESH module of the Molquest v4.5 software package (http://www.softberry.com) and Augustus was employed and 51,737 genes are predicted in the finished metassembly. Predicted numbers of genes are less compared to A1 and higher than A2.

**Table 2:** Results of Gene finding.

| **Parameter** | **A1 Assembly** | **A2 Assembly** | **A3 Assembly** |
| --- | --- | --- | --- |
| No. of genes predicted | 56,888 | 48,680 | 51,737 |

In the total gene component predicted we found 1,303 disease resistance related genes in pigeonpea. Finished assembly, A3 yielded 51,737 total genes which are less to A1 but more than reported in A2 assembly, improvement in read mapping depth results in reduction of number of earlier reported incomplete genes and yields a complete gene set. Of the predicted gene set 54-resistance single copy gene analogues containing known conserved domain NBS LRR were selected and *in silico* mapped on to the corresponding chromosomes (**Supplementary Table 2**).

### Identification of repetitive sequences and transposable elements in the improved assembly

Repeat elements are some extra copies of DNA sequences generated and planted at various locations in the genome to meet certain challenges and improve fitness during the course of evolution. Repetitive elements in Pigeonpea occupy nearly half of the genome of *Cajanus cajan* [18]. Repeats pose many computational challenges in read alignment and assembly [19] such as creation of gaps, overlaps and leading to many mapping inaccuracies in misassemblies [20]. One can always filter and exclude the reads but it is essential to map them on to chromosomal locations where gaps exist. Mate-pair libraries were used for resolving repeat problems and obtaining contiguous scaffolds in both prokaryotic [21] and eukaryotic organisms [22]. Metassembler searches for contigs that can be placed in the gap using mate pairs, and then again looks to see if there is a recorded shortest path exists between any of these contigs. In assembly, over lapping reads are used as edges to connect reads belonging to same region of genome. However in complex genomes like Pigeonpea abundance of repeats cause coverage gaps and read errors thus leaving numerous gaps to fill between contigs while scaffolding. Filling of gaps requires adoption of robust computational approaches to affectively address repeats problem. Sequenced pair-end and mate-pair reads can potentially bridge over gaps efficiently to order and orientate contigs by estimating the gap lengths to the edges while filling the scaffolding graph [23].

High level of assembly was achieved using mate-pair reads in wheat, a genome ridden with large content of repeats [24]. We analyzed the repeat content in comparison to A1 and A2 assemblies and classified them in to various classes (**Table 3**). In course of iterative use of reads during assembly we observed transposon derived repeats collapse against identical reads resulting in closure of a significant portion of gaps. Similar observations were reported on gap filling using retro transposon related repeats in human genome assembly [25].

**Table 3:** Repetitive sequences of draft assemblies A1, A2 and finished A3 assembly.

| Transposable Elements | A1 Assembly | A2 Assembly | A3 Assembly |
| --- | --- | --- | --- |
| Retrotransposons | 77,096,057 | 116,194,477 | 89,089,240 |
| Gypsy | 52,354,920 | 71,402,096 | 59,247,991 |
| Copia | 19,937,308 | 37,676,825 | 24,339,237 |
| Line | 5,261,337 | 6,717,918 | 5,914,324 |
| Unclassified elements | 216,262,607 | 169,378,278 | 158,228,382 |
| DNA transposons | 9,772,250 | 27,455,193 | 19,826,943 |
| Total transposable elements | 303,130,914 | 313,027,948 | 267,144,565 |

### Identification of microsatellites

Improved Pigeonpea assembly was mined for single sequence repeats and out of 2,98,732, 2,97,294 were simple and the remaining 1,438 of complex types. Mononucleotide repeats were the abundant with 56.05% of total SSRs, mined. Dinucleotides occupying 33.45% dinucleotides (99949), 8.72% (26069), trinucleotides and 1.27% tetranucleotides (3811) repeats. The remaining SSRs were a complex type, with 0.25% of hexa nucleotides and 0.22% of penta.

Among 167,465 mononucleotide repeats, the mononucleotide motifs were in majority with A/T repeats of 98.25% and of with the rest of. 1.74% occupied by C/G types. Among 99,949 dinucleotides microsatellites, AT/AT type (77.34%) of microsatellites were most common type in the genome followed by AG/CT type (13.21%), and AC/GT type (9.40%). The CG/CG type dinucleotides microsatellites were present at a very low proportion (0.03%). In trinucleotide SSRs repeats (26,069), around 66.71%, 12.31%, 8.07%, 5.98% of SSRs were of AAT/AAT, AAG/CTT, ATC/ATG and AAC/GTT types, and were most abundant respectively. Among the other types of repeats, the ACG/CGT type was lowest (0.36%) in the genome of Pigeonpea. The highest distribution (68.06%) of tetra nucleotides microsatellites was present in the genome of Pigeonpea. Maximum numbers of predominant SSRs repeats were of A/T type followed by AT/AT, AG/CT, AAG/CTT, AAT/ATT and AAAT/ATTT (**Supplementary Table 3**). The overall analysis showed that the relative abundance of tetra, penta and hexa SSRs types were low as compared to mono, di and tri SSR types in Pigeonpea genome sequences (**Figure 1**).

**Table 4:** Results of microsatellite search in the improved pigeonpea assembly A3.

|  |  |
| --- | --- |
| Total number of sequences examined | 13,102 |
| Total size of examined sequences (bp) | 584435790 |
| Total number of identified SSRs | 298732 |
| Number of SSR containing sequences | 6494 |
| Number of sequences containing more than 1 SSR | 4603 |
| Number of SSRs present in compound formation | 41002 |

**Figure 1:**
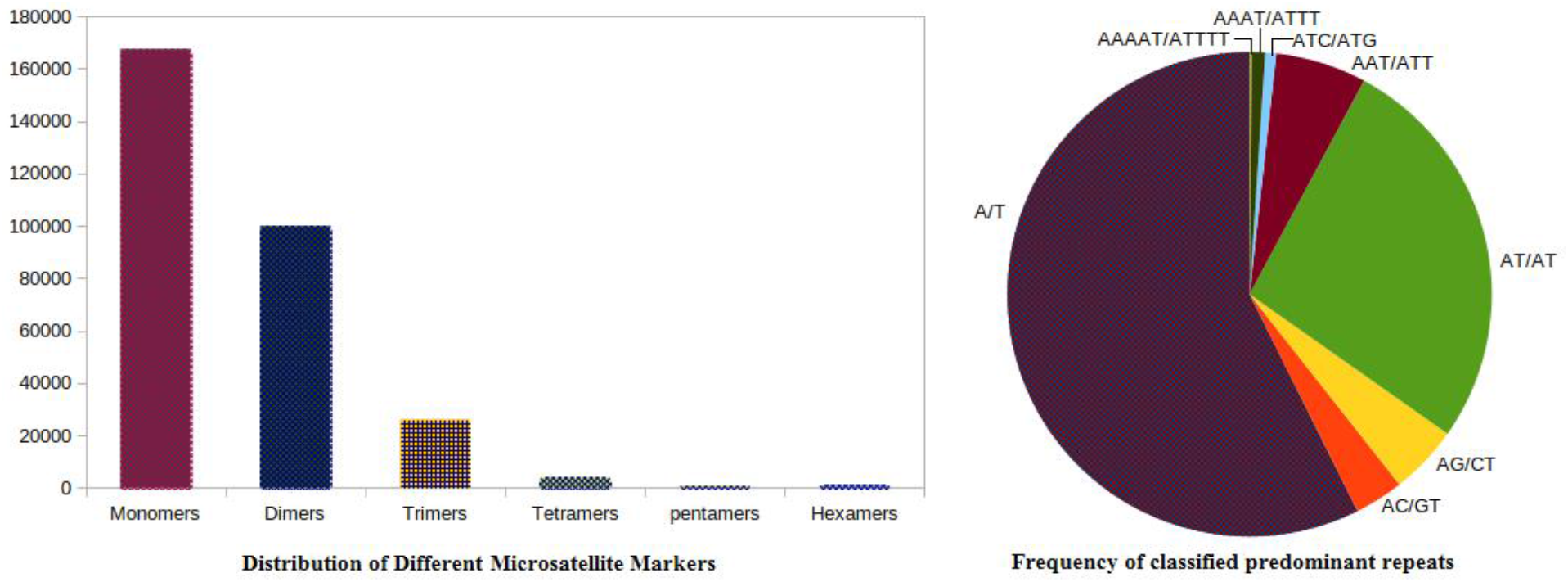
SSR distribution frequency. (A) Distribution of different repeats type classes (B) Frequency of classified predominant repeats.

### Characterization and syntenic analysis of pigeonpea NBS-LRR like resistance gene analogs

We verified the presence of already known conserved disease resistance genes in the refined metassembly. The nucleotide-binding site (NBS)-leucine rich repeat (LRR) protein sequences for other genomes were downloaded from Phytozome [26]. Comparison of predicted coding sequences against bean (*Phaseolus vulgaris*) cluster resulted in more than 100 resistance gene analogues (RGA). The predicted gene annotation revealed presence of known disease resistance domains such as ARC-NBS-LRR, Transmembrane and Kinases. Nucleotide-binding site (NBS) disease resistance genes play an important role in defending plants from a variety of pathogens and insect pests. Many R-genes have been identified in various plant species. However, little is known about presence of NBS-encoding genes in pigeonpea genome. In this study, using computational analysis of the refined genome, we identified 54 NBS-encoding single copy genes and characterized them on the basis of structural diversity and conserved protein motifs. The RGAs had high amino acid identity (77-98%) with putative disease resistance proteins in *Glycine max* several sequences with high similarity to NBS-LRR resistance (R) proteins were identified. We mined 1,301 resistance gene analogues sharing up to 78% of homology with Soybean, Chickpea, barrel clover, field bean and other species (**Supplementary Table 4**). Of them 251 NBS-LRR domain containing resistance gene analogues to pigeonpea were found (**Supplementary Table 5**).

Syntenic relationship with selected legume genomes *Glycin max* and *Medicago truncatula* revealed extensive conservation among pigeonpea and other legume plants, with 89–91 per cent of the pigeonpea assembly showing signs of RGA conservation. 41 NBS-LRR orthologs *Glycin max*, 73 NBS-LRR orthologs *Medicago truncatula*, for some 57 per cent NBS-LRR pigeonpea genes, were identified for the closely related organisms. *Glycin max* was found to have the largest number of extended conserved syntenic blocks indicating its recent ancestry followed by *Medicago truncatula*. The genome assembly of pigeonpea comprises 251 homologs of the disease resistance gene, of which 229 are anchored in pseudomolecules. The number of 41 pigeonpea genes had significant sequence homology with Glycin genes and 73 with Medicago genes. Homologous blocks containing more than 4 R genes in *C. cajan* with *G. max* and *M. trancatulum* are noted. Of these, there are 23 genes between the pigeonpea and Glycin genome assemblies with 57 collinear blocks (**Figure 2**). Overall all pigeonpea RGAs displayed extensive collinearity with different chromosomes of Glycin and Medicago. Homologous blocks connecting chr4 in *C. cajan* with chr4 of *G. max*; chr11 of *C. cajan* with chr20 and chr17 of *G. max*; Chr3 of *C. cajan* with chr19 in *G. max.* Similarly comparative analysis of draft assembly A2 [2] reported homologous blocks connecting chr3 in *C. cajan* with chr19 of *G. max*.

**Figure 2:**
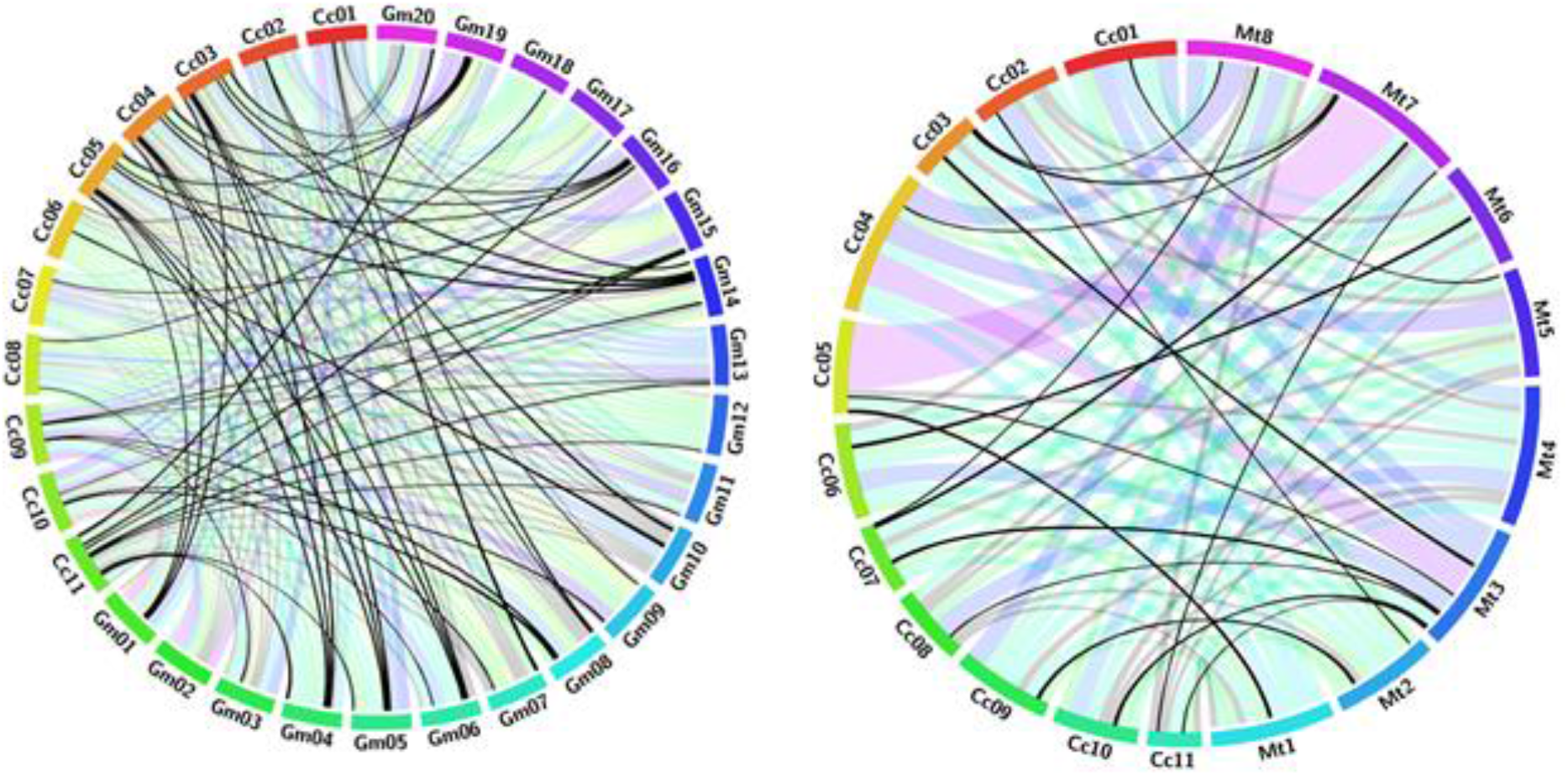
Circos diagram presenting syntenic relationship between NBS-LRR proteins from pigeonpea (Cc), *Glycin max* (Gm) and *Medicago truncatula* (Mt) pseudomolecules. Pseudomolecules of the two target species were labeled as Gm01-20 and Mt1–8. Pigeonpea pseudomolecules are labeled in different colours and labeled as Cc01-11. Colinear blocks are coloured according to the colour of the corresponding Pigeonpea pseudomolecule. Each ribbon radiating black from a pigeonpea pseudomolecule represents a NBS-LRR similarity block between pigeonpea and other legumes.

### Cloning, isolation and PCR amplification of identified putative R gene analogs (RGAs)

For designing primer sets for PCR amplification of predicted resistance gene (R) orthologs BLASTN was employed in comparison with Soybean genome. Primer sets for PCR amplification were designed using EPrimer tool [27]. PCR amplicons were eluted and sequenced by Sanger sequencing method. Isolated Pigeonpea resistance gene analogues were deposited to NCBI (**Supplementary Table 6**). List of primer sequences used in PCR amplication are given in (**Supplementary Table 7**).

Genomic DNA from 15 day old seedlings of 34 Pigeonpea cultivars was extracted employing CTAB method [28]. Purity and concentration of DNA was estimated with Nanodrops ND-1000. Nine primers were selected for polymorphism study **(Supplementary Table 7**). Polymerase chain reaction (PCR) was performed in a total volume of 20 μl containing 60 ng of template DNA, 200 μM of dNTPs, 2.5 mM MgCl2, 1x PCR buffer, 0.4 μM of each primer, 0.75 U Taq DNA polymerase and water to make the final volume up to 20 μl.

Amplification were carried out using thermocycler Bioer Gene Pro and PCR conditions was set as initial denaturation at 94°C for 5 minutes, 30 cycles of denaturation at 94°C for 30 seconds, primer annealing at 50°C for 30 seconds, primer extension at 72°C for 2 minutes and final extension step at 72°C for 7 minutes. The amplified products were visualized by ethidium bromide stained 1.5 % agarose gels in SYNGENE G-Box gel documentation unit (**Figure 3**).

**Figure 3:**
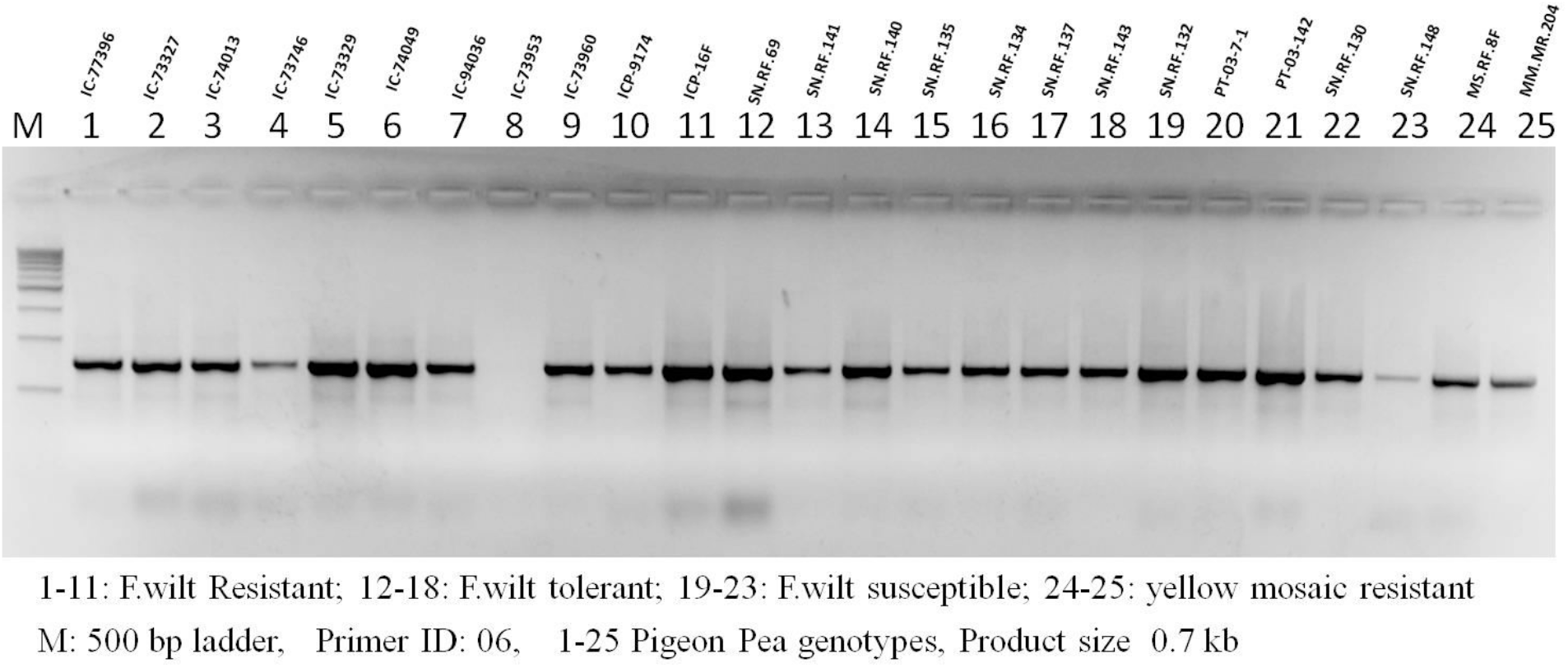
PCR amplification of *Fusarium* wilts resistant RGA among Pigeonpea genotypes.

## Discussion

In the present work we chose two available incomplete draft assemblies and employed reconciliation algorithm to correct errors. Two compared draft assemblies A1 and A2 had low genome coverage and several repeats and gaps resulting disjoin between contigs to yield lengthy scaffolds with correct contiguity. Assembly tool, Metassembler employed in the present work is based on genome reconciliation algorithm. The computational framework includes merger of two draft assemblies, A1 and A2, align them by selecting matches and mis-matches present in both, resolving gaps and other sequence errors to obtain a consensus and complete assembly.

To begin with we wanted to select the order in which the input draft assemblies are to be merged to gain subsequent superior alignment and read mapping. After several permutations, we observed that treating assembly A1 as master and aligning it with assembly A2 yielded better read mapping and lengthier scaffolds of 592,970,700 mb. Merging the two draft assemblies, in course of alignment, Meta assembler yielded matched and mismatched portions in the merged assembly by identifying homologous genomic regions with shared set of reads. Mis matches include gaps that are to be filled with right sized read sequences.

Metassembler initially utilized all available raw reads from both draft assemblies using conventional read overlapping technique to fill the existing gaps and join the contigs. However, no notable success was observed in gap filling and repeat resolution. Alternatively, we employed local pigeonpea pair-end and mate pair libraries to fill the gaps. Metassembler generated statistics, compared the distances between the mapped mates and the required sizes of insert reads to fill a gap. For example, gaps measuring < 500 mb were filled by pair-end reads while mate-pair reads were maximum utilized for filling larger gaps measuring 3 to 5 KB. Similar reports using large sized mate-pairs for filling bigger gaps in assembly of large genomes were reported [29]. In the present study employed pair-end and mate-pair reads contributed significantly to fill the gaps and thereby in joining the contigs in to full length scaffolds. Further, iterative use of pair-end and mate-pair libraries during successive alignments resulted in identification of maximal portions shared by same library of reads. This in turn has contributed to dramatic improvement of genome coverage in the resultant assembly A3. Resulting A3 assembly quality was judged using metrics-contig number, scaffold lengths, N50 and L50, genome coverage of 160x with a similarity of 72.7 %. Genome similarity score can also be useful in estimating extent of redundancy present in both genomes [30–31].

Draft assembly A1 had 360,028 contigs with N50 and L50 of 5,341 and 30,054 respectively. Reported genome coverage was 199x with a similarity of 75.6 %. Draft Assembly A2 had 72,923 contigs with N50 and L50 of 22,480 and 7,254 respectively. A2 had 592,970,700 scaffolds with reported genome coverage of 160x with a similarity of 72.7 %.

FGENESH predicted 51,737 genes using the finished metassembly, A3. Predicted number of genes are less in our finished assembly, A3 are less compared to A1 but higher than A2 (**Table 2**). Annotation of improved assembly yielded 51,737 genes predicted. Wet lab PCR amplification is the Gold standard for verification of predicted gene presence and their functionality. For PCR based gene amplification 23 primer sets were designed to screen 34 pigeon cultivars. Out of the 34 genotypes screened 14 were found to be *Fusarium* wilt resistant (**Supplementary Table 8**), 7, *F*. wilt tolerant, 5 *F*. wilt susceptible, 5 yellow mosaic susceptible genotypes (**Figure 3**). Data on yellow mosaic disease reaction is not presented here. PCR amplified genes were isolated, cloned and submitted to NCBI (**Supplementary Table 6**). Genotype, environment interaction in the field determines the phenotypic performance of isolated plant genes [32]. Phenotypic evaluation of predicted resistance genes in field trials is also required for transfer of the obtained results to pigeonpea downstream breeding programs for development of disease resistant cultivars. Field experiments were conducted to assess the disease reaction of predicted R genes to *Fusarium* wilt taking cv. Asha (object of present study) as control with 34 Pigeonpea cultivars. The replicated field experiments were conducted at Ranchi (Jharkhand state) and Rahuri (Maharastra), India during 2011, 2012 and 2012, 2013 rainy seasons. Of the 26 screened cultivars against check cv. Asha, 14 resistant and 6 tolerant at Ranchi farm and at Rahouhuri farm 8 resistant, one tolerant and 6 susceptible disease reaction was observed to the F. wilt disease of Pigeonpea. Observed variation in disease incidence reflect the natural agro climatic conditions prevailing at the individual trail site.

## Conclusion

In the present work genome reconciliation algorithm was adopted to reconstruct draft assemblies to produce an accurate and near complete genome assembly of pigeonpea. We demonstrated successful implementation of our reassembly frame work by merging two chosen draft assemblies employing pair-end and mate-pair libraries to correct gaps and other sequencing errors. Resulting reconstructed metassembly was superior to compared two draft assemblies in terms of measured assembly quality statistics *viz.* N50 and scaffold lengths. Quality of finished assembly was assessed for presence known conserved resistance gene loci (imparting resistance to *Fusarium* wilt disease in Pigeonpea). Annotation of improved assembly yielded prediction of 1303 resistance genes (including six extra genes gained from metassembly). PCR screens and field experiments validated the resistance reaction of isolated genes against *Fusarium* wilt thus making the results available to Pigeonpea breeders.

## Methods

We developed a workflow model (**Figure 4**) based on reconciliation algorithm, that includes 1. Merging two mis-assemblies, 2. Finding matches and mismatches and other sequencing errors, 3. Closing gaps using pair-ends, mate-pair libraries, 4. Assessment of finished assembly quality, prediction of disease resistance gene families, their isolation and characterization.

**Figure 4:**
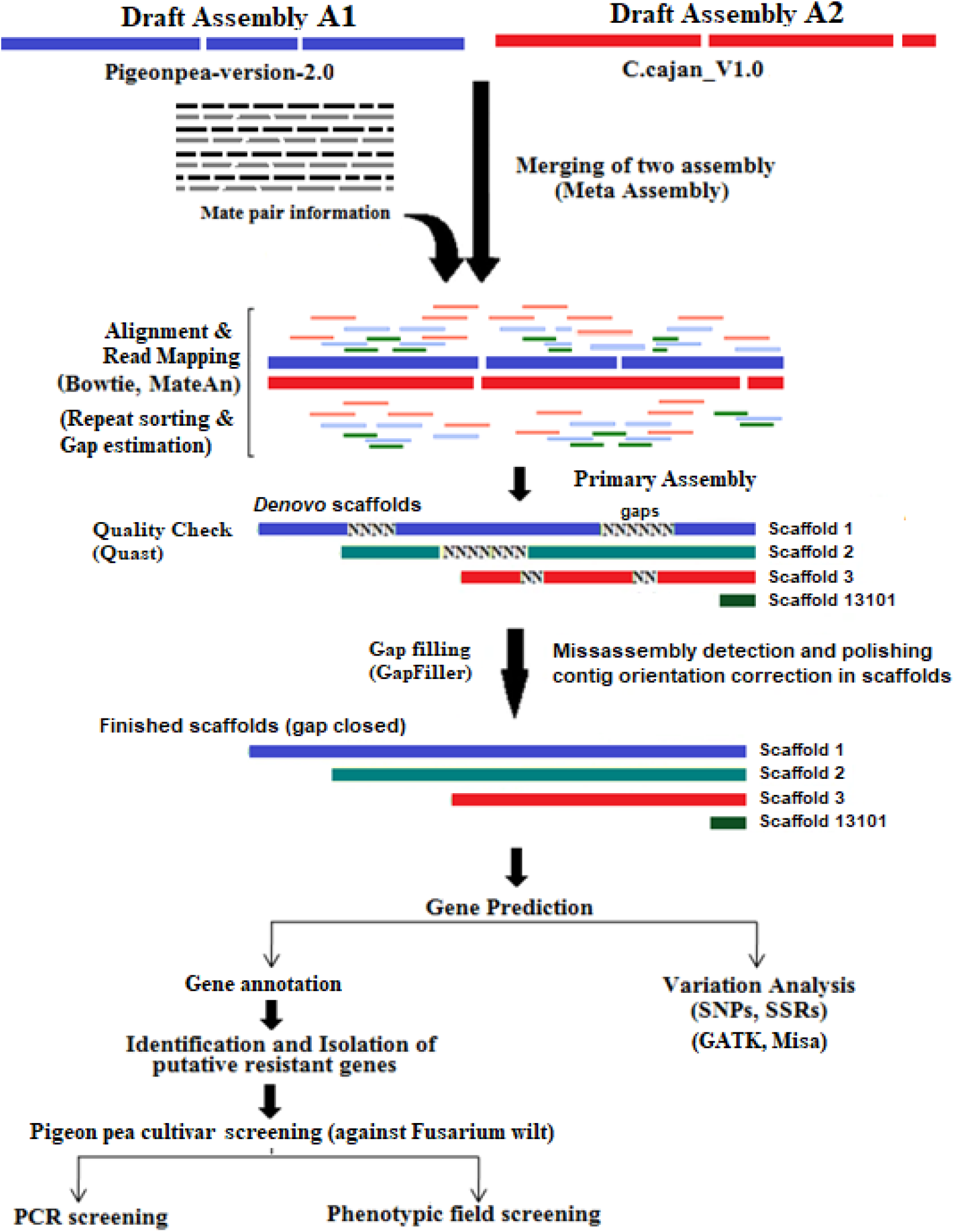
Experimental Frame work depicting reconstruction steps of Pigeonpea genome.

### Retrieval of pigeonpea genome datasets

Complete data sets belonging to two whole genome sequences of Pigeonpea and associated 23 SRA reads were downloaded from the National Center for Biotechnology Information (NCBI) (https://www.ncbi.nlm.nih.gov) to the local storage-GCA_000340665.1 (SRA accessions SRR5922904-SRR5922907) and GCA_000230855.2 (SRA accessions SRR6189003-SRR6189021) for the *cv* Asha.

### Genome reconstruction and quality assessment

Illumina pair-end and mate-pair library sequence reads of Pigeonpea, *cv* Asha were quality checked using FASTQC v0.11.8 (http://www.bioinformatics.babraham.ac.uk/projects/fastqc). Contaminated reads were removed to get error corrected reads. Reads with sequence quality of Phred scores of less than Q30 (base calling accuracy with less than 99.99%) were removed using PRINSEQ v0.20.4 (https://sourceforge.net/projects/prinseq) and reads were repaired using BBmap v37.66 (https://sourceforge.net/projects/bbmap).

Reported Pigeonpea draft assemblies A1 [1] and A2 [2] were both sequenced employing Illumina technology and assembled with SoapDenovo v2.3.1 assembler. In present work, data sets A1 (GCA_000340665.1 consisting 4 SRA read sets) and A2 (GCA_000230855.2 of 19 SRA read sets) were analyzed employing reconciliation algorithm [11]. The work flow includes the steps: 1) Merger of two mis-assemblies, 2) Finding matches, mismatches and other structural errors, 3) Closing gaps using pair-end, mate pair libraries, 4) Assessment of finished assembly quality 5) Prediction of disease resistance gene families, their isolation and characterization. A1 consisted of 360,028 initial contigs (N50 5341, 648 Mb) with 30% of gaps within contigs. A2 contained 72,923 scaffolds (N50 22480, 605 Mb) with 20% of intra scaffold gaps. We used all the read datasets available belonging to A1 and A2 with NCBI. All the computations including read pre-processing, quality control, comparison of two draft assemblies, their alignment, gap filling, assembly merger, map accuracy, quality assessment, putative gene prediction were performed on HPC server employing Meta Assembler [33].

GapFiller [34] was employed to find the existing gaps (A1 30%; A2 20%). Initially short reads were used for filling gaps, resulting A1 genome size of 648 Mb and 605 MB of A2 draft assembly.

We initially employed overlap approach with available read sequences followed by used pair-end as well as mate-pair library data sets for resolving repeat redundancy, gap filling and other structural errors. Firstly, we used entire single paired read data sets available (minus two mate-pair data sets) of A1 along with all data sets from A2. Alternatively, second treatment included two mate-pair data sets from A1 and all full data available from A2. At the end of analysis, all the output values and statistical metric data were collected for comparative performance analysis.

Draft assembly A1 was sequenced in 2011 and had genome coverage of 199% [1]. However, using the same raw read data, the authors had again reassembled employing A3 assembler and reported gain of coverage, i.e. an increase of ~15% (from 60.0% to 75.6%,) and resubmitted to NCBI. In our present work we used this recent assembly set, A1 along with A2 assembly data [2] for reassembly and improvement (**Figure 4**).

We observed that in our reassembly pair-end insert read sizes below 500 bp in our library were maximum utilized for filling smaller gaps. Mate-pair sizes up to 5.0 kilo base pairs are available in our library. In our metassembly these mate pairs were employed affectively used for closing medium and long distanced gaps (even up to 20-25 kb). Similar results on use of large sized mate-pairs for filling bigger gaps was reported in assembly of large genomes [29].

### Merging misassemblies and gap closure

Draft assembly sequences A1 and A2 were merged in to a single sequence. Alignment and merger of A1 and A2 assemblies resulted in a total scaffold length of 548 Mb. Resulting merged assembly is compared to A1 and A2 draft assemblies (75.6% and 72.7% respectively) has an improved genome coverage of 82.4 %. Yet the Merged sequence contained 10% of gaps.

To improve further contiguity and accuracy of merged sequence existing intra scaffold gaps were filled. Repeat content and existing gaps were estimated employing Gapcloser and Gapfiller tools [34]. In the second round of gap filling various computational approaches such as paired end, mate pair libraries and remaining unused short reads were used. Gap content, estimation of repeats (**Table1**). Iterative use of left over short reads (300bp) contributed to filling nearly 20% of gaps. After polishing and another round of reassembly yielded a scaffold length 13,348 (scaffolds of N50 574,622) with a coverage of 174x %.

### Finished genome assembly and quality assessment

Increased N50, maximum scaffold length and minimum number of contigs, increased N50 values together with longer scaffolds contribute to improved genome coverages. In mis-assemblies the number of gaps and ‘N’s caused due to repeats were measured. In course of metassembly we strived to minimize gaps and other sequencing errors. We employed Quast v4.5 [35] to gather extensive assembly statistics. BUSCO v3.2 [17] was employed for assessing the genome completeness, annotation and sets of predicted genes. Mapping accuracy and identification of resistant gene analogue loci were assessed. In addition 75% of unigenes were aligned to the reassembled genome.

### Gene prediction and function annotation

Metassembly was first repeat-masked using RepeatModler and Repeat Masker tools [36], followed by *ab initio* gene prediction using the FGENESH module of the Molquest v4.5 software package (http://www.softberry.com). The predicted genes were annotated using BLASTX (E<10^6^) search against the NCBI non-redundant (nr) protein database using Blast2GO software [37]. Synteny blocks between the genomes of pigeonpea and other legumes were computed by blastp combined with the Circos [38] to understand homology to the NBS-LRR gene from *Glycin max* (Gm) and *Medicago truncatula* (Mt) pseudomolecules.

### Identification of genome wide SSR

Refined genome sequence of Pigeonpea was analyzed identify to various Single Sequence Repeat markers (SSRs) types using Microsatellite Identification tool (MISA) (http://pgrc.ipkgatersleben.de/misa/). Minimum length for SSR motifs per unit size was set to 10 for mono, 6 for di and 5 for a tri, tetra, penta, hexa motifs. We calculated the total lengths of all mono-, di-, tri-, tetra-, penta-, and hexa-nucleotide repeats in terms of base pairs of SSR per mega base pair (Mb) of DNA.

### Gene validation

Genome similarity score recorded set of sequenced reads originating from one draft genome correctly mapped on to a second genome. To check the accuracy in finished Pigeonpea genome we wanted to verify the location of certain genomic regions or loci present in the inputted two assemblies. A set of genes imparting resistance against various pests and diseases are located in B4 cluster on chromosomes in two examined draft assemblies of pigeonpea, (*Cajanus cajan*) Asha. As a test case location of B4 gene cluster syntenic regions was verified in the present study to estimate the accuracy of read mapping achieved in the finished assembly.

### Computational resources

We run all reassembly and merging using HPC Cluster having CentOS-Linux version 7,2.93 GHz 2x Intel Xeon 8 core processors and 2 TB of RAM. Majority of the running time is spent on assembly process and about 1/4 on graph construction and analysis. However, Reconciliator uses more than 1.5 TB of RAM to merge the Asha isolates, Pigeonpea assemblies.

## Data availability

The improved draft genome assembly of Pigeonpea is available at NCBI/ENA/GenBank, under the Accession Number WWND00000000.

## Acknowledgments

Authors thank ICAR-National Bureau of Plant Genetic Resources, New Delhi for providing research facilities, and Centre for Agricultural Bioinformatics (CABIN, ICAR-IASRI), New Delhi India for providing high performance computing (HPC) facility.

## Contributions

SM conceptualized and supervised all the experiments, interpreted the results and wrote the manuscript text. PM and RM performed all the high-throughput bioinformatics analysis, software and formulated the manuscript. MS conducted field phenotypic experiments, SNR and RK updated the final manuscript for publication. DPW, AKG and SKS performed all the wet lab experiments.

## Supplementary Information legends

**Supplementary Table 1:** BUSCO (Benchmarking Universal Single-Copy Orthologs) genes distribution.

**Supplementary Table 2:** Putative Disease resistance genes predicted from improved reassembly of Pigeonpea.

**Supplementary Table 3:** Numbers of predominant SSRs repeats.

**Supplementary Table 4:** Resistance gene analogues sharing homology with species.

**Supplementary Table 5:** NBS-LRR domain containing resistance gene analogues to Pigeonpea [*Cajanus cajan* (L.) Millsp.].

**Supplementary Table 6:** List of Pigeonpea disease resistance genes submitted to NCBI.

**Supplementary File 7:** List of primer sequences used in PCR amplication.

**Supplementary Table 8:** Pigeonpea phenotypic field scores for *Fusarium* wilt disease reaction.

